# Leveraging multiplicity in biologically informed neural networks to uncover disease heterogeneity

**DOI:** 10.64898/2026.07.15.738681

**Authors:** Dennis Gankin, Pedro Beltrao

## Abstract

Biologically inspired neural networks (BINNs) embed pathway, ontology, or protein-interaction structure directly into neural networks, promising interpretable disease prediction where hidden nodes map to named biological entities. Yet BINNs have been hard to train at biobank scale, and the reliability of their interpretations remains largely untested. Here we present a fast BINN implementation trained on UK Biobank genotype and plasma proteomics data from about 500,000 individuals across six common diseases. BINNs achieve competitive predictive performance, but we uncover two major limits to their interpretability. First, attribution scores are strongly biased by graph topology, because node degree and layer position influence the scores. Normalization reduces this bias but can weaken enrichment for known disease genes. Second, BINNs show substantial predictive multiplicity, that is, independently trained models with identical architecture and data reach similarly accurate solutions while prioritizing different genes and pathways. Although this multiplicity makes single-model explanations unstable, the range of interpretations can itself reveal disease biology. Across 100 replicate BINNs for type 2 diabetes, we find distinct solution clusters prioritizing either inflammatory or hepatic-metabolic pathways, mirroring known disease heterogeneity. Thus, analyzing the space of BINN explanations can turn multiplicity into a tool for studying complex disease mechanisms.

## Introduction

Deep learning is increasingly used in biology, advancing diverse areas of biological research. Models such as AlphaFold have revolutionized protein structure prediction and opened new avenues for biological discovery (Jumper et al. 2021; Lin et al. 2023). In parallel, sequence models have emerged as powerful tools for biological sequence analysis and enable accurate variant effect predictions ranging from clinical consequences to molecular functions and regulatory mechanisms (Cheng et al. 2023; Avsec et al. 2026, 2021). More recently, foundation models for protein sequences, DNA, and single-cell data provide new ways to represent and learn from unlabeled datasets, creating opportunities for generative modeling and downstream prediction tasks (Hayes et al. 2025; Brixi et al. 2025; Theodoris et al. 2023).

Despite their predictive power, deep learning models often remain black boxes because they provide limited insight into the internal logic underlying their predictions. However, in biology, deep learning approaches are increasingly used not only for prediction but also for explanation, making model interpretability essential. Models trained on genomic, proteomic, or clinical data are often used post hoc to nominate disease genes, prioritize pathways, and generate hypotheses about biological mechanisms (Eraslan et al. 2019; Novakovsky et al. 2023). Interpretability becomes especially important for understanding complex human traits or diseases, where many molecular and cellular processes can contribute to the same outcome (Lappalainen et al. 2024).

Most interpretability approaches explain trained models post hoc, often by analyzing internal weights, estimating feature importance, or performing input perturbations. In sequence models these methods can help identify input patterns driving the predictions, for example, Min et al. (Min et al. 2017) analyze convolutional kernel activations across sequence inputs to uncover sequence motifs relevant for transcription factor binding site prediction. More recently, researchers have also applied sparse autoencoders and concept learning methods to extract higher level features, such as concepts used by a vision transformer for cancer prediction from histopathology slides (Kim et al. 2026).

Explaining a single trained model, however, assumes that its explanation is representative of the patterns supported by the data. Yet similar predictive performance can arise from different learned features, internal representations, and sample-level predictions, a phenomenon known as predictive multiplicity (Marx et al. 2020; Watson-Daniels et al. 2023). As a result, two similarly performing models may therefore yield substantially different explanations (Fisher et al. 2019). While predictive multiplicity has been studied more broadly, its implications for interpretability in biological deep learning remain largely unexplored. Biologically informed neural networks (BINNs) are particularly well suited for studying interpretive multiplicity because their biologically structured architecture allows explanations to be compared across independently trained models in terms of genes and higher-order functional annotations (Yu et al. 2018). This is especially relevant for disease risk prediction from genotype or proteomics measurements, where black box deep learning models can achieve strong predictive performance but typically provide limited mechanistic insight into disease biology (Selby, Sprang, et al. 2025; Rudin 2019).

Since DCell (Ma et al. 2018), such visible or knowledge-primed neural networks have been applied to various biomedical prediction tasks, including cancer drug response (Kuenzi et al. 2020), prostate cancer classification (Elmarakeby et al. 2021), single-cell analysis (Fortelny and Bock 2020), often producing competitive predictive results and plausible biological interpretations. Their appeal is that trained models can be interpreted directly in terms of relevant genes, pathways, and biological processes. However, recent studies and reviews have cautioned that initialization, training choices, and biases in the underlying biological knowledge graphs can influence the interpretations (Esser-Skala and Fortelny 2023; Selby, Jakhmola, et al. 2025). While existing work mostly reports single-model explanations or average explanations across a small number of runs, the broader distribution of interpretations has not been studied. At the same time, applying BINNs to biobank-scale cohorts and high-dimensional genomic inputs remains computationally challenging, so previous UK Biobank applications have relied on reduced input representations or smaller sample sizes (Lee et al. 2024; van Hilten et al. 2021).

Here, we systematically characterize interpretive multiplicity in biologically informed neural networks trained at biobank scale. We introduce FastBINN, a faster BINN implementation that enables training on UK Biobank-scale cohorts of approximately 500,000 individuals and makes large-scale replicate experiments tractable. Using FastBINN, we train disease-specific genotype based BINNs for six common conditions and examine biases and instability of the interpretations, and we ask whether the distribution of explanations can itself be biologically informative. We find that raw attributions are strongly confounded by graph structure and require normalization. More importantly, we show that interpretive multiplicity persists even across similarly predictive models and after reducing feature redundancy with proteomic inputs. In type 2 diabetes, 101 replicate proteomics based models reveal a structured, low-dimensional interpretation space with two dominant explanatory modes: one enriched for inflammatory pathways and another for metabolic and liver-related biology. These results suggest that multiplicity in BINNs is not merely implementation noise, but may reflect the mechanistic heterogeneity of complex traits. More broadly, we propose using the solution space of well-performing models as a tool for interpretation and discovery, helping to identify distinct biological mechanisms that may contribute to complex traits.

## Results

### Scalable Biologically Informed Neural Networks enable interpretable complex trait prediction at biobank scale

Biologically informed neural networks (BINNs) provide mechanistic explanations of their predictions and could therefore be useful for investigating the biological processes underlying disease risk (Selby, Sprang, et al. 2025). Here, we present FastBINN, an optimized implementation that scales BINNs to large biobank cohorts for genotype analyses of complex human traits. FastBINN builds neural networks in which input features are connected through prior biological knowledge rather than dense, unconstrained layers. In genotype based models, we first map approximately 700,000 single nucleotide polymorphisms (SNPs) to genes. Then, we map the genes to Gene Ontology annotations using either the biological process or cellular component graph. We train a separate model for each disease trait to learn trait-specific predictive signals. We also apply the same framework to proteomic input data, where the smaller feature space of 2,707 measured proteins makes it feasible to train large sets of replicate models and analyze the explanation space in greater detail. Because predictions pass through annotated biological units, we use Integrated Gradients (Sundararajan et al. 2017) to attribute trait-specific predictive signals to genes, biological processes or cellular components at population scale (Fig. 1a).

**Figure 1:**
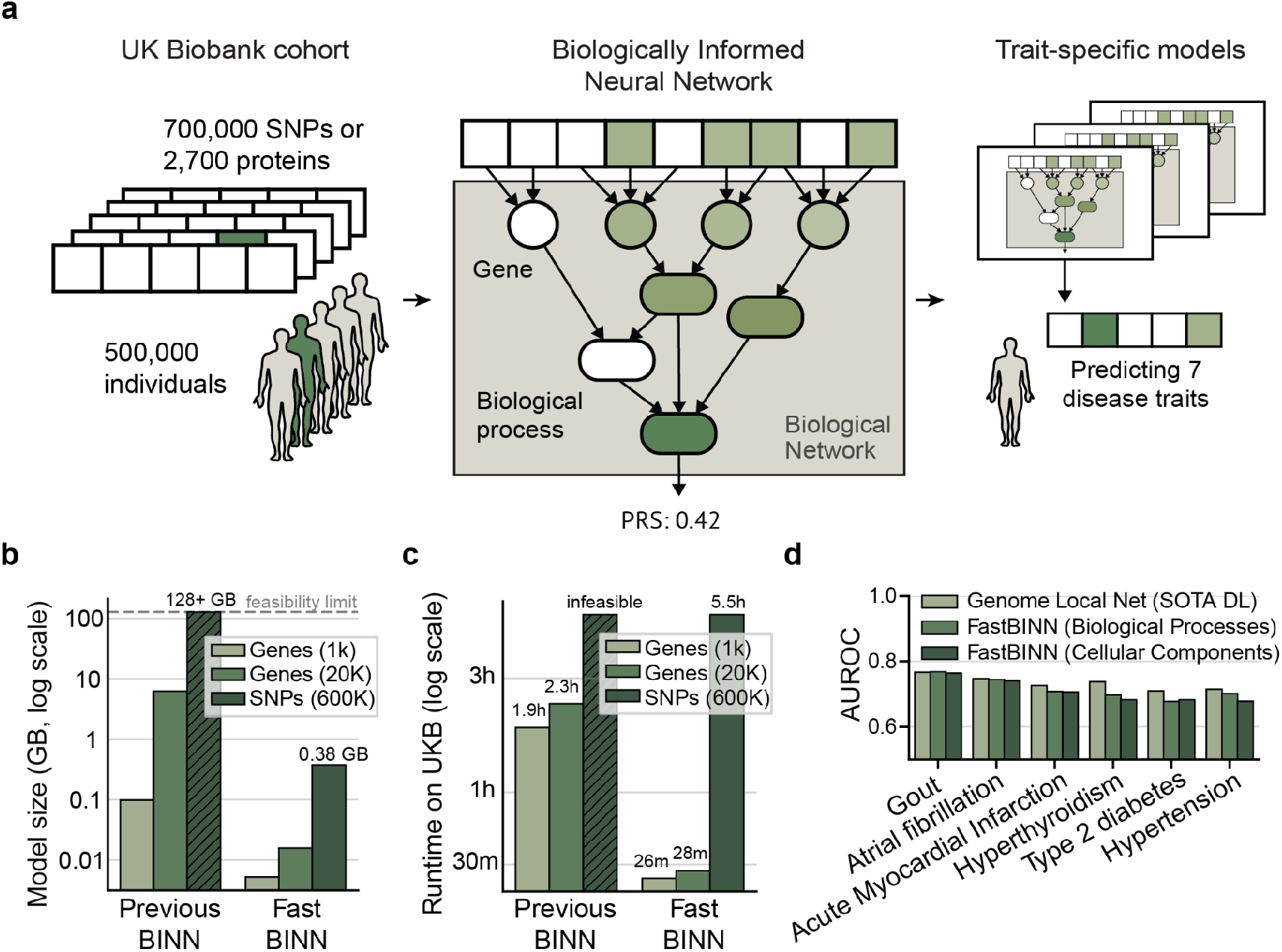
FastBINN enables interpretable disease-risk prediction at UK Biobank scale. **a.** Overview of the FastBINN framework. Genotype input features are connected to genes and then to higher-order Gene Ontology annotations. Separate models are trained for each disease trait. After training, attribution scores link predictive signal to genes and ontology terms. **b.** Memory requirements for different model configurations. The previous BINN implementation (Park et al. 2024) exceeds the 128 GB memory limit for genome-wide SNP inputs, whereas FastBINN reduces the model size sufficiently to load it into memory on single compute nodes. **c.** Runtime comparison between a previous BINN implementation from Park et al. 2024 and FastBINN across input scales. FastBINN reduces training time over 10 epochs by more than two orders of magnitude, enabling full genome-wide SNP models to be trained within 6 hours. **d.** Predictive performance across seven common disease traits. FastBINN models based on biological process and cellular component graphs are compared with Genome Local Net, an unconstrained deep learning benchmark for genotype based prediction (Sigurdsson et al. 2023). FastBINN approaches benchmark performance for several traits, with biological process models generally outperforming the more constrained cellular component models.

Applying such models to large cohorts, such as the UK Biobank, has so far been computationally challenging. The high dimensionality of genome-wide genotype data creates large networks with many input-layer connections, and scaling these models to hundreds of thousands of individuals substantially increases memory requirements, computational complexity, and training time. As a result, training BINNs on full genotype-scale data has remained largely intractable. In practical terms, for about 700,000 input SNPs a previous BINN implementation from Park et al. 2024 exceeds the 128 GB memory limit of the available large GPU compute nodes of the UK Biobank computational cluster. FastBINN addresses this bottleneck through memory-efficient graph construction and model representation, reducing the model size for genome-wide SNP inputs to 0.38 GB and enabling full-scale biologically structured neural networks to be loaded into memory on single UK Biobank compute nodes (Fig. 1b).

FastBINN also markedly reduces training time. Through optimized computation and matrix operations, runtimes decreased by over 100-fold from multi-hour training with a previous implementation (Park et al. 2024) to minutes for proteomics input and 5.5 h for genome-wide SNP models on UK Biobank data (Fig. 1c). These improvements enable us to train BINNs on full UK Biobank genotype data across multiple common disease traits, rather than restricting analyses to reduced feature sets, simplified networks or smaller cohorts. Moreover, in the smaller proteomics setting, they also make it feasible to train large sets of replicate models, allowing us to systematically analyze variability in model explanations.

The trained genotype models approached the performance of Genome Local Net, a state-of-the-art black box deep learning model for genotype based prediction (Sigurdsson et al. 2023). Prediction performance is closely matched for several traits, including gout and atrial fibrillation, whereas larger gaps remained for others, including type 2 diabetes and hypertension (Fig. 1d). So for some traits, performance may be limited by the sparsity of the ontology-based architecture, which restricts the model to effects represented in current annotations. Consistent with this possibility, models based on the larger biological process graph tended to perform slightly better than the more constrained cellular component models (Fig. 1d). Together, these results show that FastBINN enables full-scale, interpretable neural prediction of complex disease traits while retaining competitive performance against unconstrained deep learning models.

### Attribution scores recover known disease biology but have network bias

The main advantage of BINNs over black box Deep Learning models is their network’s interpretability by design (Selby, Sprang, et al. 2025). Hence, applying them to complex traits we set out to investigate the biological relevance of the resulting model interpretations. After training a BINN, we compute attribution scores of genes and Gene Ontology terms to the prediction outcome. This yields an importance metric for all biological units in the network in the context of the prediction. To assess the biological relevance of the model attributions of our genotype based model, we compared highly attributed genes with known disease-associated genes. For all predicted traits, known disease genes were overrepresented among the top-ranked genes by the raw attribution score. This overrepresentation increases toward the top of the ranking and among the top 0.5% most highly attributed genes, fold enrichment ranged from 13.9 for hyperthyroidism to 1.4 for atrial fibrillation. These differences may reflect variation in disease complexity and in the number of known disease-associated genes available for each trait (Fig. 2a). While the prioritization of disease genes indicates that the attribution scores are biologically meaningful, they exhibit strong correlation with the topology of the neural network graph (Fig. 2b). That is because, as reported in earlier studies (Esser-Skala and Fortelny 2023; Fortelny and Bock 2020), the underlying neural network architecture biases the attribution scores. Specifically, the neural network layer a node is in, dominates the node’s attribution score, with a spearman correlation of -0.9 between attribution score and node layer. This is an effect of the gradient based attribution scoring, as the gradient decreases with deeper layers in neural networks. In addition to this, central nodes in the neural network with high degree or closeness centrality exhibit higher attribution scores. Thus, the raw attribution scores are network-biased and not fully disease-specific, which confounds gene prioritization and the biological interpretation of the predictions. Consequently previous works have pointed out the importance of normalizing these scores to remove network bias (Esser-Skala and Fortelny 2023).

**Figure 2.**
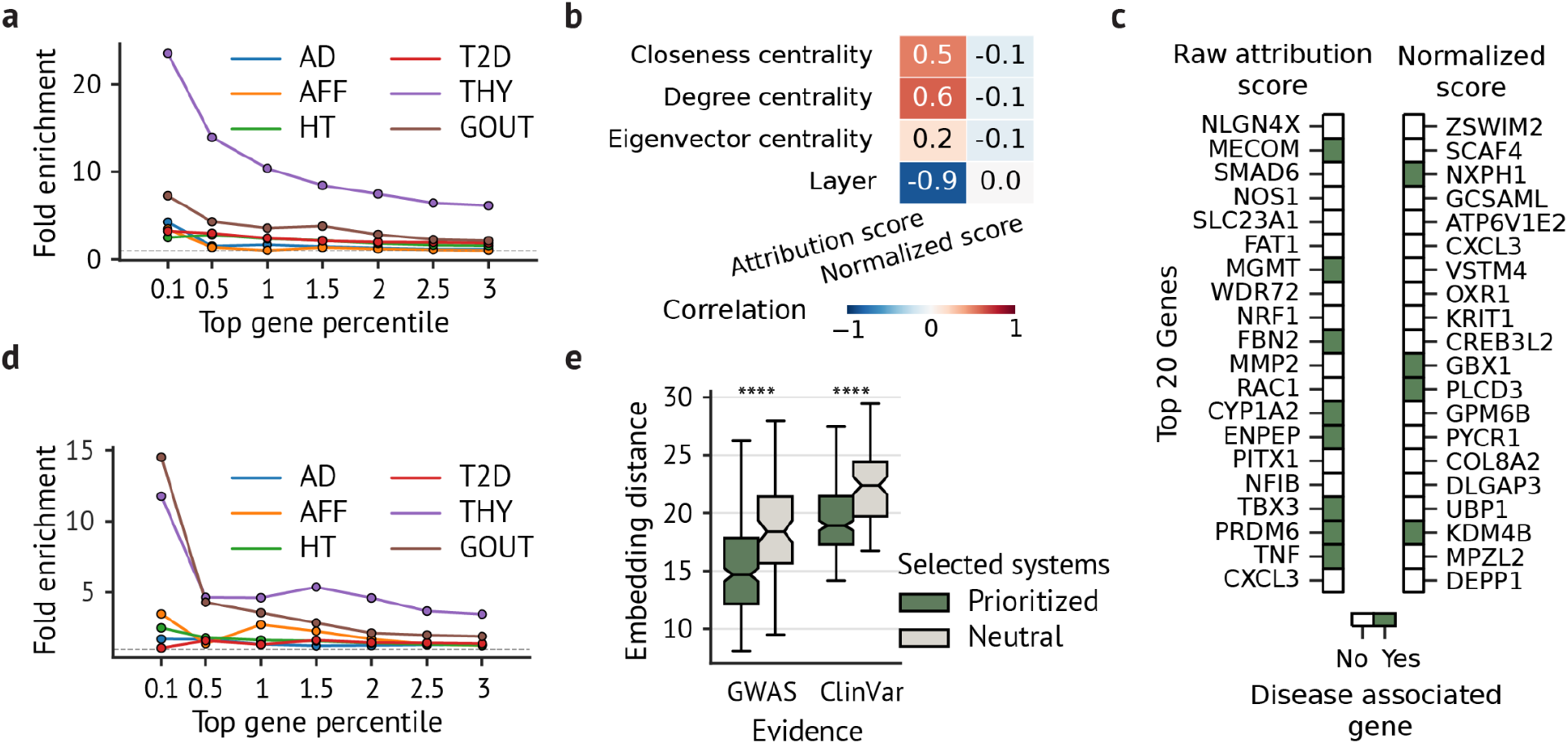
FastBINN attributions recover disease-relevant genes after correcting for network bias. **a.** Raw attribution scores recover known disease-associated genes across traits, with strongest enrichment among the highest ranked genes. Fold change enrichment shown for models trained to predict Alzheimer’s disease (AD), type 2 diabetes (T2D), atrial fibrillation (AFF), hyperthyroidism (THY), hypertension (HT), and gout (GOUT). **b.** Correlation between attribution scores and network properties. Raw attribution scores correlate with graph properties such as centrality and layer position, whereas normalized scores show reduced dependence on these features. **c.** Example of the effect of normalization on gene prioritization. The top-ranked genes can change substantially after correcting attribution scores. **d.** Fold enrichment of known disease-associated genes after normalization of attribution scores. Normalization reduces enrichment relative to raw scores but preserves disease relevant signal above background. **e.** Independent validation of normalized FastBINN attributions using a latent-neighborhood prioritization framework. Biological processes prioritized by FastBINN are closer in embedding space to disease representations from Open Targets (GWAS) and the European Variation Archive (ClinVar) than size-matched sets of mid-ranked biological processes sampled around the 50th percentile of the normalized attribution scores, supporting the biological relevance of normalized attributions beyond direct GWAS overlap.

Normalizing against 100 models trained on randomly shuffled labels considerably reduces the network bias and lowers the correlation of attribution scores with network metrics (Fig. 2b), making the attributions more disease-specific (Fig. S3). At the same time, the normalization substantially changes the gene importance rankings. Thus, gene prioritization based on normalized attribution scores can return very different top genes compared to raw attribution scores that are not corrected for network bias (Fig. 2c). However, the normalization also reduced the fold enrichment of disease-associated genes among highly attributed genes for most traits. In hyperthyroidism, enrichment among the top 0.5% of most highly attributed genes decreased from 15-fold to 5-fold. Nevertheless, the normalized scores still enrich disease associated genes indicating that the models capture biologically relevant signals beyond what is explained by the network topology (Fig. 2d).

To further validate the biological relevance of the normalized FastBINN attribution scores, we compared them with an orthogonal disease gene prioritization method based on network propagation (Trip et al. 2026). This method embeds genes, traits and biological terms in a shared biological space, integrating external disease gene evidence from diverse genetic and functional evidence sources. Related diseases and terms occupy nearby regions of this space, allowing disease neighborhoods to prioritize candidate genes using information not used to train FastBINN. Biological processes prioritized by FastBINN were significantly closer to the corresponding disease embeddings from genome-wide association study (GWAS) evidence and rare variant evidence (ClinVar), compared to equally sized sets of non-prioritized biological processes (FDR adjusted p = 2.6e-9; Fig. 2e). While the GWAS evidence used to construct these embeddings may partly include data from the same cohort used to train FastBINN, the ClinVar evidence provides independent rare-variant input to the embedding model and therefore offers an orthogonal validation. So, even though network topology affects raw FastBINN attributions, normalized attributions retain disease-relevant biological signal supported by independent data.

### Predictive multiplicity induces interpretive multiplicity in BINNs

While normalization can control for network bias and still yield relevant interpretations, another reported caveat in BINN interpretability is the robustness of explanations across independent training runs with different seeds (Esser-Skala and Fortelny 2023; Fortelny and Bock 2020). To feasibly study this on a large scale, we turn to smaller and better performing proteomics models, that is BINNs trained to predict traits from plasma protein abundance measurements of 2,707 proteins. In this analysis, we focus on type 2 diabetes prediction, as it is a common trait. The blood protein levels in humans provide a reliable readout of human health, so the proteomic BINNs demonstrate a much higher AUROCs for type 2 diabetes than the genotype based models (AUROC 0.68 vs. 0.89; Fig. 3a). Yet, as predictive models, BINNs exhibit the effect of predictive multiplicity. Hence, two training runs with different initialization seeds can reach similar prediction performance of 0.86 and 0.88 AUROC respectively but yield relatively different predictions across samples (Spearman rho=0.45; Fig. 3b).

**Figure 3.**
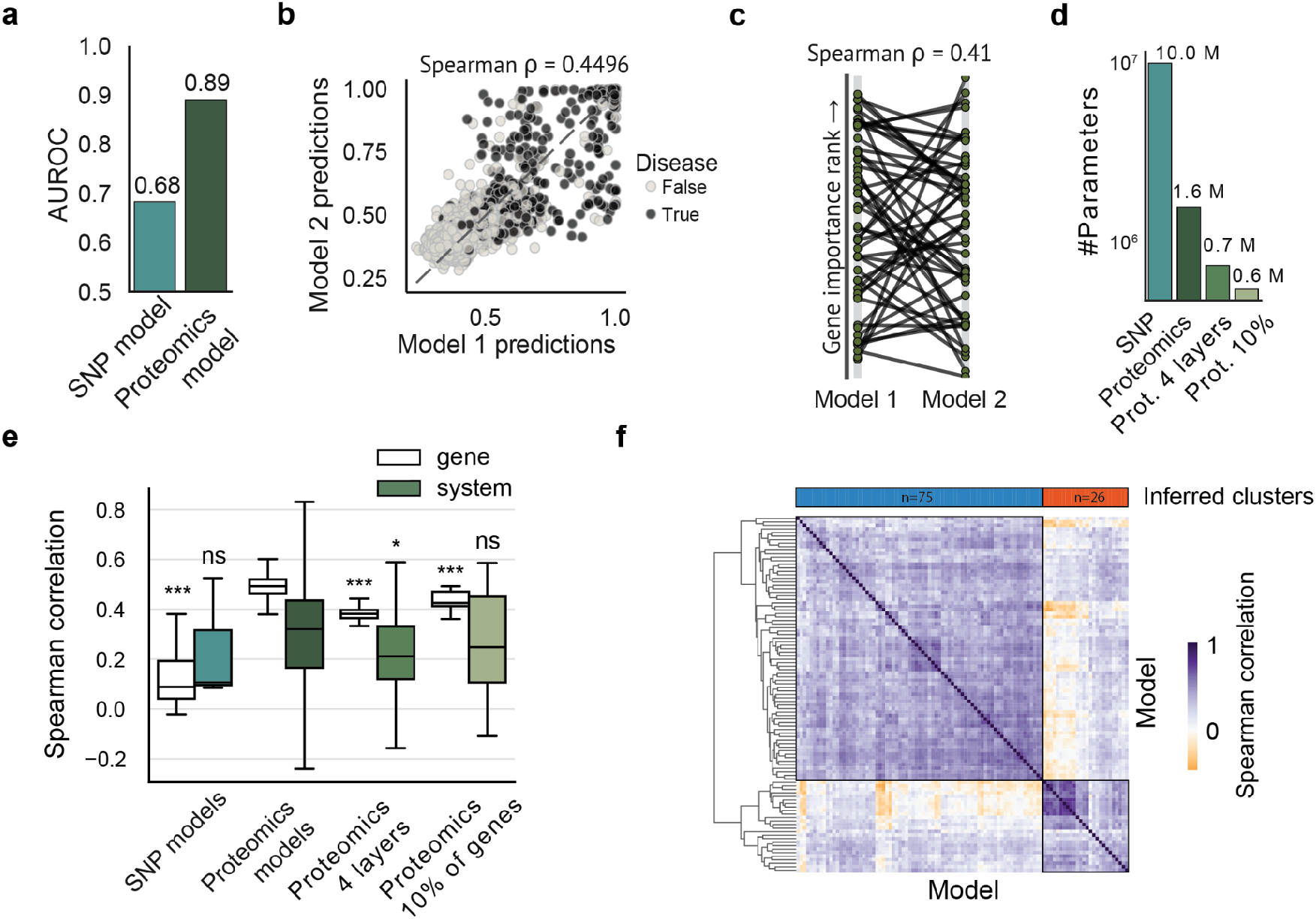
Predictive and interpretative multiplicity persists across FastBINN models. **a.** Predictive performance of SNP based and proteomics based FastBINN models. **b.** Example of predictive multiplicity between two proteomics based FastBINN models trained with different random seeds. Both models achieve similar prediction performance, but their predictions per sample deviate. Points indicate individuals and are coloured by disease status. **c.** Example of interpretative multiplicity between the same two models. Gene importance ranks differ substantially across two runs, showing that similarly predictive models can prioritize different genes. **d.** Number of trainable parameters across model configurations. Proteomics based models are substantially smaller than SNP based models, and additional simplifications further reduce model size. **e.** Pairwise Spearman correlations of gene- and system-level attribution profiles across independently trained models of different input types and sizes. Proteomics models show higher agreement than SNP based models on average, but correlations remain variable, particularly for higher-order biological systems. Significance is compared to the proteomics model. **f.** Correlation structure across 101 trained proteomics FastBINN models with similar predictive performance on type 2 diabetes. Hierarchical clustering of attribution profiles reveals distinct groups of models, indicating structured regions in the explanation space.

This predictive multiplicity has implications for model interpretation, too. As the models can reach the same prediction quality through different parameter sets, their resulting explanations will also vary, leading to interpretative multiplicity (Marx et al. 2020). Studies in various application fields show that models trained with different random seeds yield different model explanations (Pawelczyk et al. 2020; Paulo and Belrose 2025; Hwang et al. 2026). FastBINN also demonstrates such interpretative multiplicity where the attribution scores of two individually trained models demonstrate a Spearman correlation of 0.41, as illustrated by Fig. 3c. The stark difference in the gene importance rankings between the two models indicates that models that learn to predict different risk values for individuals, then also diverge on their feature interpretations. So similar to findings of unstable BINN interpretations (Esser-Skala and Fortelny 2023; Fortelny and Bock 2020), attribution scores of FastBINN models trained on human disease prediction also vary across training runs.

As shown above, model multiplicity demonstrates strong influence on the downstream interpretation, but the extent of this effect and its driving parameters in BINNs remain unclear. Hence, we wanted to further explore the interpretive multiplicity to see how it changes with model size, input dimensionality and prediction performance. Moving from genotype to proteomics input for this analysis reduced the input dimensionality from about 700,000 input SNPs to 2,700 protein levels, decreasing the FastBINN model size from 10M to 1.6M parameters (Fig. 3d). To assess whether input redundancy or network depth contributed to multiplicity, we constructed two additional model variants. The first used only the 10% most informative proteins as input to reduce redundancy, whereas the second pruned the network to a maximum depth of four layers. These changes partially reduce interpretive multiplicity. On average, the attribution scores agree between proteomics based replicate models, demonstrating spearman correlations of 0.5 for gene-level attributions and 0.3 for system-level attributions. The correlation of gene-level attribution scores is therefore significantly higher in proteomics based models than in the genotype based models (p=2.8e-13; Fig. 3e). This suggests that higher prediction performance and smaller model size influence the average agreement between model interpretations. Yet, the higher system-level attribution agreement for proteomics based models compared to genotype based models is not statistically significant. Also, the depth-reduced and redundancy-reduced proteomics models do not significantly impact the correlation of system-level attribution scores between replicates. That is likely due to the large variance of system-level correlations across the proteomics model pairs, ranging from highly correlated pairs above 0.8 to slightly anti-correlated pairs below −0.2. So, while higher prediction performance and a smaller network can lead to more robust interpretations on average, individual replicate pairs can still be very different. Thus, with such a large space of explanations, interpreting two randomly picked models could yield very different biological hypotheses.

To explore the interpretation space in more detail, we analyse 101 proteomics models predicting type 2 diabetes with comparable performance. The pairwise correlation of the system attributions between the training runs reveals a clear structure in the explanation space. Some clusters of models have highly correlating interpretations with each other but strongly differ from other model interpretations, with two main explanation clusters standing out (Fig. 3e). This indicates that model explanations occupy distinct regions of interpretation space, rather than simply showing stochastic variation. Consequently, considering only a single trained model or a simple average of model explanations would obscure this structure. Therefore, we asked if the variability between similarly predictive models might encode biologically meaningful alternative representations of type 2 diabetes risk.

### The interpretation space of equally predictive BINNs reveals alternative biological mechanisms

Seeing the structure in the interpretation space of the 101 trained proteomics models for type 2 diabetes, we investigated if this variation contains biologically meaningful insights into disease mechanisms. First, we validated the model attributions by comparing it to an orthogonal data integration approach that creates a shared embedding space of related genes, biological terms and diseases from genetic evidence (Trip et al. 2026). Although the 101 proteomics based models prioritized different biological processes, the processes prioritized across FastBINN models are significantly closer in the embedding space to type 2 diabetes than non-prioritized processes (Fig. 4a). This demonstrates that the attribution scores of biological processes in FastBINN models capture biologically relevant information.

**Figure 4.**
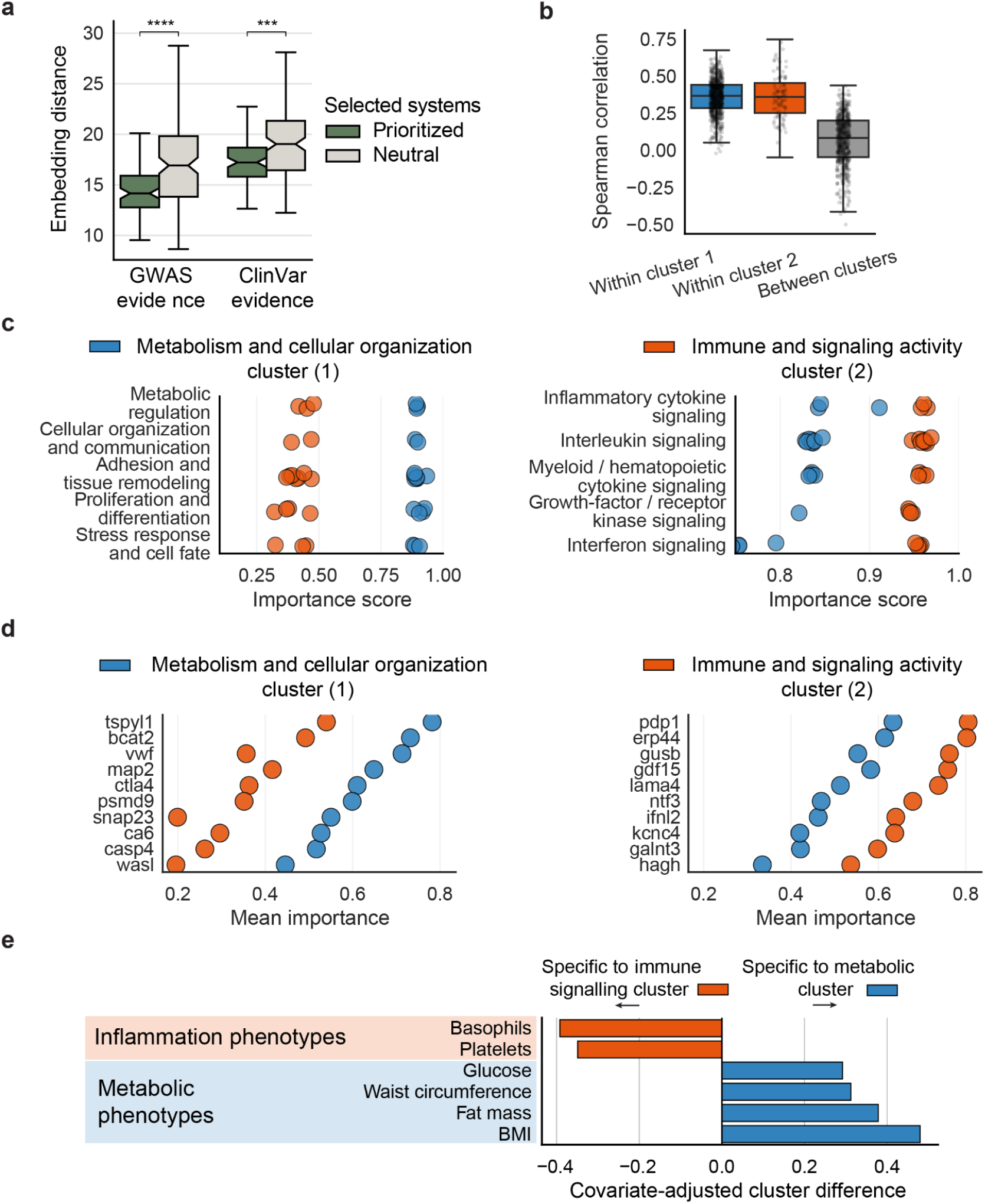
Multiplicity reveals distinct biological explanations for type 2 diabetes prediction. **a.** Independent validation of biological processes prioritized by FastBINN using a latent-neighborhood data integration framework. Prioritized biological processes show smaller embedding distances to disease representations from Open Targets (GWAS) and the European Variation Archive (ClinVar) than size-matched sets of mid-ranked biological processes sampled around the 50th percentile of the normalized attribution scores, supporting the biological relevance of the processes prioritized by the proteomics based models. **b.** Pairwise Spearman correlations of attribution profiles across independently trained models. Correlations are higher within the two major model clusters than between clusters, showing that replicate models form distinct groups with internally more consistent explanations. **c.** Biological processes preferentially prioritized by the two major clusters. One cluster emphasizes metabolism and cellular organization, the other focuses on interleukin and cytokine signalling. **d.** Genes with the largest cluster-specific differences in attribution. The metabolism and cellular-organization cluster and the immune and signalling cluster prioritize distinct sets of genes in line with the biological processes, indicating that cluster-level differences extend from system to gene-level explanations. **e.** Phenotypic differences of highly predicted individuals between the two model clusters. Bars show the covariate-adjusted difference in each phenotype. High risk individuals predicted by the metabolism-associated cluster show higher metabolic phenotypes, while platelets and basophils are elevated in individuals prioritized by the immune signalling cluster.

The biological signal captured by the FastBINN models varies between models. Model explanations formed two main clusters of interpretations, with stronger attribution score correlations within clusters than between them. The broad spread of correlations suggests additional fine-grained structure within the interpretation space of each cluster (Fig. 4b and Fig. 3e). Together with the validated biological relevance of the prioritized processes, this structured interpretation space suggests that different explanation clusters may capture alternative biological signals related to type 2 diabetes. Investigating this space further through biological processes with high, cluster-specific attribution scores, revealed that the two explanation clusters prioritize different biological themes. One cluster was primarily associated with metabolic, structural, and tissue-remodeling related processes, including cellular organization, adhesion, proliferation, and metabolic regulation, while the other prioritized immune, cytokine, and signaling-related processes, including interleukin, interferon, and growth factor signaling pathways (Fig. 4c). This separation is consistent with the current understanding of type 2 diabetes as a complex disease with both metabolic dysfunction and inflammatory or immune-mediated components (Alfadul et al. 2022).

The divergence of explanations also extended to the gene level, where the most different genes prioritized by the two model groups reflect the distinct biological mechanisms (Fig. 4d). The cluster associated with metabolic and structural processes prioritized genes linked to metabolic regulation, vesicle trafficking, cellular organization, and vascular or structural functions, including *BCAT2*, *SNAP23*, *VWF*, and *CTLA4*. In contrast, the immune signaling-associated cluster prioritized genes connected to inflammatory signaling, extracellular matrix remodeling, cellular stress responses, and interferon-related activity, including *GDF15*, *LAMA4*, *IFI44*, and *XIAP*. Together, these findings suggest that different groups of models converge on biologically meaningful yet distinct mechanisms underlying type 2 diabetes prediction.

The split of biological themes between the model clusters also reflects how the trait manifests in different individuals. Models in the metabolism and cellular organization associated cluster assign a higher disease risk to individuals with high metabolic phenotypes, such as waist circumference and fat mass (Fig. 4e). In contrast, models in the inflammation and signalling related cluster predict higher risk scores for individuals with high platelet and basophil counts, two indirect indicators of inflammation. These findings indicate that the space of model interpretations may correspond to the heterogeneity of manifestations or progressions of type 2 diabetes in individuals. So interpretive multiplicity of a predictive model alone can be biologically meaningful and analysing the interpretation space can provide a richer picture of the complex disease mechanisms underlying the trait prediction.

## Discussion

Independent validation of normalized FastBINN attributions using a latent-neighborhood disease gene prioritization framework. Biological processes prioritized by FastBINN are closer in embedding space to disease representations from Open Targets (GWAS) and the European Variation Archive (ClinVar) than size-matched sets of mid-ranked biological processes sampled around the 50th percentile of the normalized attribution scores, supporting the biological relevance of normalized attributions beyond direct GWAS overlap.

In this study we show that analyzing interpretative multiplicity in biologically informed neural networks can reveal alternative biological mechanisms of complex diseases. As an interpretable-by-design model for complex trait prediction, we present FastBINN, a fast biologically informed neural network scaled to biobank-scale genotype data. FastBINN enables large-scale disease prediction from genotype or proteomics data with integrated gene- and process-level interpretability, and thus makes it feasible to explore the explanation space of multiple independently trained models. In line with previous studies, our results show that the topology of the underlying biological knowledge graph can strongly bias model attributions (Esser-Skala and Fortelny 2023; Fortelny and Bock 2020). In BINNs, central or highly connected nodes receive higher raw attribution scores, reflecting graph structure in addition to predictive signal. Part of this topology may reflect real biological organization, such as broadly acting processes. However, Gene Ontology and other knowledge graphs also include strong non-biological biases arising from incomplete, uneven annotations skewed towards well-studied processes and genes. Less biased architectures could be constructed from clustered large-scale protein-interaction data (Laman Trip et al. 2026), but at the cost of direct interpretability of the created clusters. Normalization reduces topology-driven attribution bias, but it may also remove biological signal encoded in the structure of the network itself.

We demonstrate that predictive multiplicity occurs in FastBINN models, with similarly performing models assigning very different disease risk scores to the same individual. Thus, the same individual may be classified as high risk by one model but low risk by another, complicating the use of such models in medical and health applications. However, understanding this variability may also guide ensemble strategies by identifying distinct solutions that contribute complementary information. The predictive multiplicity also extended to model interpretation, where similarly performing models prioritize different genes and biological systems. Multiplicity remains even in smaller proteomics based models with higher prediction performance compared to genotype based models. Exploring 101 replicate model explanations for type 2 diabetes revealed a structured interpretation space, where the main explanation clusters mapped to biologically plausible mechanisms. Across both systems- and gene-level analyses, one cluster emphasized metabolic, structural, and tissue-remodeling processes, while the other prioritized inflammatory, cytokine, and immune-signalling related mechanisms, indicating that interpretive multiplicity reflects distinct disease-relevant aspects of type 2 diabetes biology.

Although predictive multiplicity weakens the robustness of interpretations of independent models, the resulting variance in model explanations can be biologically meaningful. However, our analysis provides only an initial view of interpretive multiplicity in BINNs, and several questions remain open. First, we explored the interpretive multiplicity in proteomics based models that are smaller and more predictive than genotype based models. Future work should test whether the larger and less predictive genotype based models also exhibit such structured interpretation spaces. Due to the large number of replicate models needed this is computationally more challenging.

Our results suggest that network size, and input correlation and dimensionality affect interpretive multiplicity, but we cannot yet determine how these factors shape the solution space. Other questions about factors controlling multiplicity in BINNs remain. This includes how predictive a model needs to be for interpretable explanation clusters to emerge, and how many replicate models are needed to capture the interpretation space in a biologically meaningful resolution. In addition, factors such as model architecture, sparsity, regularization and the attribution method likely influence interpretive multiplicity, too, and their effects need to be studied systematically.

Lastly, this opens up new avenues to investigate complex traits with predictive models by applying similar analyses to additional diseases, less well-studied traits and other molecular data modalities. Additionally, to reproducibly characterize, and compare interpretation spaces across traits, cohorts and data modalities, we need to establish a general analysis framework. If validated and generalized, interpretation space analysis could provide a useful approach for generating mechanistic hypotheses from complex trait prediction models.

More broadly, our findings connect to a general challenge of model interpretation under multiplicity in machine learning. Predictive multiplicity is a known effect in machine learning and often model ensembles tackle the challenge of prediction robustness (Watson-Daniels et al. 2023; Black et al. 2021). For interpretation however, multiplicity poses the challenge that similarly performing models rely on different internal representations (Marx et al. 2020). So ensembles could mask model specific signals, but explanations of a single model may provide only one view of the signals supported by the data. The problem of interpretation multiplicity and instability spans across many machine learning fields and interpretability methods. Not only attribution scores in biological deep learning models, but also in general settings counterfactual explanations and sparse autoencoders suffer from multiplicity (Paulo and Belrose 2025; Hwang et al. 2026; Pawelczyk et al. 2020; Esser-Skala and Fortelny 2023). Our results suggest that interpretation variability should not only be treated as a source of instability, but can also be analyzed as a structured space of alternative explanations. Studying this space may help identify which signals are robust, which are model-specific, and which correspond to distinct but scientifically meaningful solutions.

Overall, this study motivates a refined view on global model interpretation under interpretive multiplicity. We show that the diversity of explanations supported by similarly accurate models can contain biological information that may be obscured by average explanations. Thus, interpretation multiplicity should not only be treated as a robustness problem, but also as a space of explanations that can be explored to uncover alternative biological mechanisms in complex traits.

## Methods

### UK Biobank data

We analyzed seven common disease traits in the UK Biobank, selected on the basis of their prevalence, known genetic contribution, prior use in genotype based prediction studies and representation in the complex-trait literature. Case and control status was assigned using ICD-10 diagnosis codes derived from UK Biobank data field 41202, resulting in prevalences shown in Supplementary Fig. S1. For genotype based models, we used UK Biobank genotyping-array data from approximately 500,000 unrelated individuals passing standard genotype quality control. Variants and individuals were filtered with PLINK v1.9 using minor allele frequency, variant missingness and individual missingness thresholds of --maf 0.001, --geno 0.03 and --mind 0.1, respectively. The SNP genotypes were one-hot encoded for input to FastBINN. Plasma proteomic measurements were obtained from UK Biobank data field 30900 for 52,634. Protein abundance values were filtered for missingness, log-transformed and batch-normalized. For the correlation reduced proteomics representation, proteins were clustered according to their abundance profile similarity, and a subset comprising 10% of proteins with the highest signal variance was retained.

### FastBINN network architecture

FastBINN models were implemented as sparse feed-forward neural networks whose connectivity follows prior biological knowledge from the Gene Ontology (GO). We used the Biological Process (BP) and Cellular Component (CC) GO branches, removed obsolete terms, and mapped genes to GO terms, using the GO release 2024-09-08 (https://release.geneontology.org/2024-09-08/index.html). Each gene or GO term was represented by five neurons across two layers, and units were connected according to the underlying GO graph, following the BINN framework described in (Park et al. 2024). For genotype based models, SNPs were one-hot encoded and mapped to genes using the Open Targets SNP-to-gene mapping pipeline downloaded on June 26, 2024. SNPs not mapped to any gene, genes without input features or GO connections, and GO terms without upstream input were removed. The resulting networks contained 695,641 input SNPs. For proteomics based models, log-transformed protein abundances were used as one-dimensional inputs and mapped to their corresponding genes based on the gene identifiers provided by the UK Biobank before propagation through the GO network, resulting in 2,707 protein abundance input values. Terms without upstream protein input were pruned. For additional proteomics model experiments, we generated a depth-reduced network by collapsing all GO terms at layer 5 or deeper into their corresponding parent terms at layer 4. This produced a smaller and more balanced architecture with a maximum GO depth of four layers.

### Improved BINN implementation and training

FastBINN models were trained as binary classifiers to predict disease risk for each trait. The model output was passed through a sigmoid activation and compared with the binary disease label using binary cross-entropy loss. We extended the BINN implementation of Nest-VNN (Park et al. 2024) to improve memory efficiency and training speed. The main changes were the use of sparse tensors across the whole model, vectorization of the SNP-to-gene input layer, and a hidden model state that is updated layer by layer. This enabled sparse layered operations on unstructured biological networks, including genotype networks in which input SNPs have variable path lengths to the output node. For model training, individuals were split into training, validation and test sets using a 70/15/15 split. Model selection was based on validation performance, and the checkpoint with the highest validation AUROC was evaluated on the held-out test set. Reported predictive performance corresponds to test-set AUROC. FastBINN performance was compared with the reported AUROC performance of Genomic Local Net, a black box genotype based deep learning benchmark (Fig. 1d). Model training was performed on the UK Biobank Research Analysis Platform using the mem2_ssd2_gpu1_x64 compute node with an NVIDIA A10G GPU, 22GB GPU memory and 250GB RAM for the genotype based models and the smaller the mem2_ssd2_gpu1_x16 compute node for the proteomics based models (NVIDIA A10G GPU, 22GB GPU memory, 64GB RAM). Genotype based models were trained for 10 epochs, as validation performance typically saturated within this range and each training run required approximately 6 hours. Hyperparameters were tuned separately for each trait using the validation set. Proteomics based model training took around 15 minutes per model with 10 epochs.

### Attribution score computation and normalization

Attribution scores were computed using Integrated Gradients (Sundararajan et al. 2017). For each biological unit, corresponding to either a gene or a Gene Ontology term, attributions were calculated with respect to the unit’s output neuron. For genotype based models, the non-mutant allele was used as the Integrated Gradients baseline. Attributions were computed for each individual and then averaged across individuals to obtain global gene- and term-level attribution scores. To correct for attribution biases induced by network topology, observed attribution scores were normalized against a null distribution obtained from 100 models trained on randomly shuffled disease labels. For proteomics based models, null distributions were generated by training models on the same input samples, but with randomly shuffled disease labels. For genotype based models, 100 null models were trained on a local compute cluster using synthetic genotype inputs sampled from the empirical mutation distribution, together with randomly shuffled labels, to approximate the full-input null distribution at feasible computational cost. Attribution scores were log-transformed, z-scored relative to the node-specific null distribution, and subsequently percentile-normalized.

### Model benchmarking (Fig. 1)

To benchmark predictive performance, we compared the best performing FastBINN model against previously reported ROAUC values for the Genomic Local Net (GLN), the current state-of-the-art deep learning approach, across seven disease traits (Sigurdsson et al. 2023). While GLN is a sparse feed-forward neural network, it contains less sparsity and is based on a much more structured network, than the GO based FastBINN. To assess scalability, we benchmarked model size and runtime using simulated input data, run locally on a single cluster compute node equipped with an NVIDIA RTX 4090 GPU and 128GB RAM. We tested 3 different input sizes (1K, 20K, 600K) mimicking proteomics and genomics input mapped onto the GO Biological Process network architecture. For the runtime analysis we assumed 500,000 samples and a training run of 10 epochs. At the largest input scale (600K SNPs), the previous BINN architecture exceeded the available system RAM (>128 GB) during model instantiation and could not be loaded. We therefore did not attempt to benchmark its runtime at this scale and report it as infeasible.

#### Attribution score validation (Fig. 2)

Gene-level attributions were validated by testing whether highly attributed genes were enriched for known disease associated genes. Disease gene associations were obtained from the Open Targets genetics portal (https://platform.opentargets.org/downloads) and the GWAS Catalog (https://www.ebi.ac.uk/gwas/). For each trait, enrichment was computed for genes in the top attribution percentiles. Fold enrichment was calculated as the proportion of known disease genes among top-ranked FastBINN genes relative to the corresponding background proportion across all genes represented in the network. Acute myocardial infarction was excluded from the analysis because fewer than ten known disease genes were available. System-level attributions were validated using an external embedding space of diseases and biological terms from (Trip et al. 2026), computed by network propagation from GWAS and ClinVar evidence. Disease identifiers and GO term identifiers were mapped to the corresponding embedding coordinates. For each trait, top-ranked GO terms were selected by normalized attribution score, and their embedding distances to the corresponding disease representation were compared with distances for an equal number of non-prioritized terms sampled from the 50th percentile of the attribution ranking. Statistical significance was assessed with a two-sided Mann-Whitney U-test and corrected for multiple testing using the Benjamini-Hochberg false discovery rate. Separate embedding distance analyses by trait are shown in Fig. S4.

#### Predictive and interpretive multiplicity analysis (Fig. 3)

To analyse the extent of predictive multiplicity we trained replicate models using the same architecture, input data and hyperparameters, but different random seeds. Due to feasibility we performed the main multiplicity experiments using proteomics based FastBINN models, which were smaller and computationally more tractable than the genotype based models. Because genotype based models were computationally expensive to train and attribute, only a small number of genotype replicates were generated. Type 2 diabetes was selected for these experiments because it is common in UK Biobank and is well predicted from plasma proteomic measurements. For the main analysis, we trained 101 proteomics based FastBINN models for type 2 diabetes. In addition, smaller sets of proteomics based models were trained for the depth-reduced and redundancy-reduced network configurations (6 input redundancy reduced models and 9 network depth capped models). For each replicate model, gene- and system-level attributions were computed as described above. Interpretive similarity between models was quantified using Spearman correlations between their attribution profiles, separately for gene-level and system-level attributions. The resulting pairwise correlation matrix was visualized as a clustered heatmap. Models were hierarchically clustered based on attribution profile similarity using the euclidean distance metric and the group-average linkage method. The two main interpretation clusters were defined by the first major split of the hierarchical clustering dendrogram.

#### Interpretation space analysis (Fig. 4)

The 101 proteomics models were assigned to the two main interpretation clusters identified by hierarchical clustering. For each cluster, we calculated the median normalized attribution score of every biological process across models in that cluster. Cluster-specific biological processes were identified by comparing mean attribution scores between clusters after choosing the 50 most important processes for each cluster. Cluster specificity was quantified as the difference in the mean normalized attribution score between the two clusters. For visualization and interpretation, the 20 most cluster-specific biological processes from each cluster were manually grouped into five broader biological categories. The complete list of cluster-specific biological processes is provided in Supplementary Fig. S6. Similarly, at the gene level, cluster-specific genes were identified by ranking genes according to the difference in mean normalized attribution scores between clusters. For each cluster we show the 10 genes with the highest absolute mean score difference between clusters. To relate the two interpretation clusters to physiological phenotypes, we tested whether the disease risk scores predicted by models in each cluster were associated with a set of inflammation- and metabolism-related traits in the UK Biobank. We derived data from the following UK Biobank data fields: 30080 (platelet counts), 30220 (basophil percentage), 48 (hip circumference), 30740 (glucose), 23100 (whole body fat mass), 21001 (body mass index). For each of the two model clusters, we selected individuals with type 2 diabetes who had strong cluster-specific predictions, defined as a predicted score in the top quartile (>= 75th percentile) of the cohort that also exceeded their predicted score for the other cluster. Each phenotype was converted to a signed robust z-score. Cluster differences between phenotypes were then assessed by ordinary least-squares regression of each phenotype’s signed z-score, adjusting for year of birth and genetic sex. The resulting coefficient quantifies the covariate-adjusted mean difference between clusters.

## Acknowledgements

This work uses data provided by patients and collected by the NHS as part of their care and support. This research has been conducted using the UK Biobank Resource under application number 129007. PB is supported by the Helmut Horten Stiftung and the ETH Zurich Foundation.

## Data availability

The UK Biobank data is available to approved researchers through the UK Biobank access procedure (https://www.ukbiobank.ac.uk/use-our-data/apply-for-access/). Gene Ontology data was downloaded from (https://release.geneontology.org/2024-09-08/index.html). The disease gene associations used in this study were obtained from Open Targets (https://ftp.ebi.ac.uk/pub/databases/opentargets/platform/24.09/output/etl/parquet/associationBy <u>DatasourceIndirect/</u>) and the GWAS Catalog (https://www.ebi.ac.uk/gwas/).

The FastBINN training and analysis code used in this work is available on GitHub (https://github.com/DennisGankin/fastbinn_pub).

## Supplements

**Figure S1.**
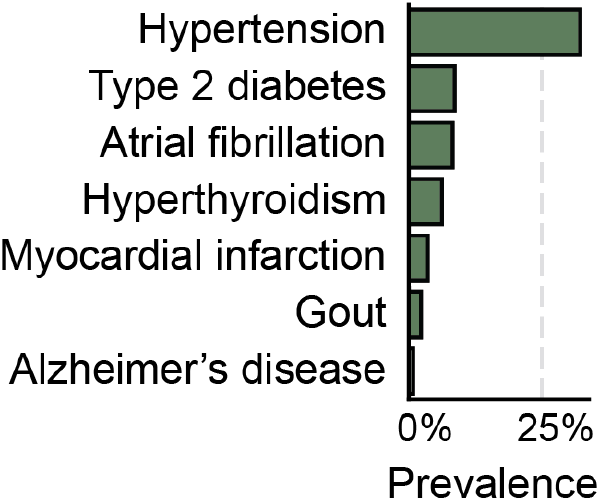
Disease prevalence in the UKB cohort of selected common diseases. Prevalence in the UK Biobank cohort of the seven common diseases the FastBINN genomics model was trained on.

**Figure S2.**
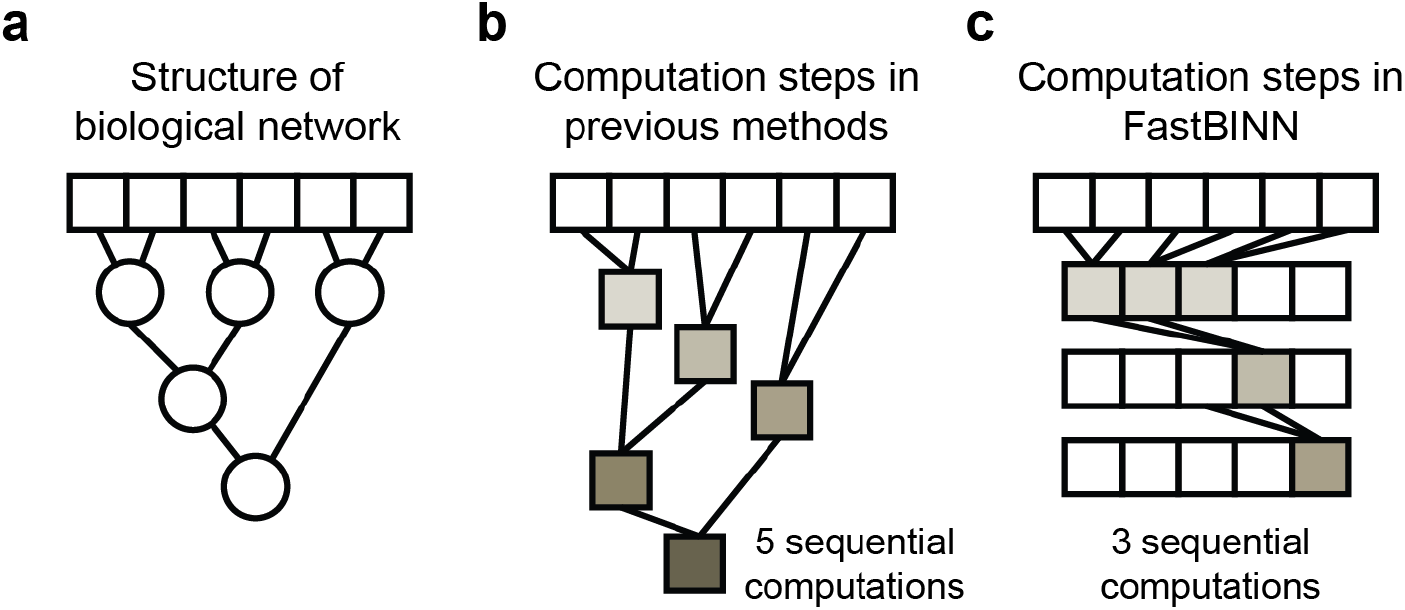
Layered computation in FastBINN reduces the number of sequential operations on unstructured biological networks. **a.** Structure of a biological network, in which input features (squares, e.g., SNPs) connect through intermediate biological entities (circles, e.g., genes and pathways) to a single output node. The biological is a directed acyclic graph but does not have a layered structure, as in classic feed-forward neural networks. **b.** Previous BINN methods propagate information node by node along these paths, so the number of sequential computations scales with the number of nodes in the network. **c.** FastBINN instead assigns nodes to layers and maintains a single hidden model state that is updated one layer at a time, enabling sparse layered operations over the hierarchical, but unstructured network. This significantly reduces the number of sequential computations.

**Figure S3.**
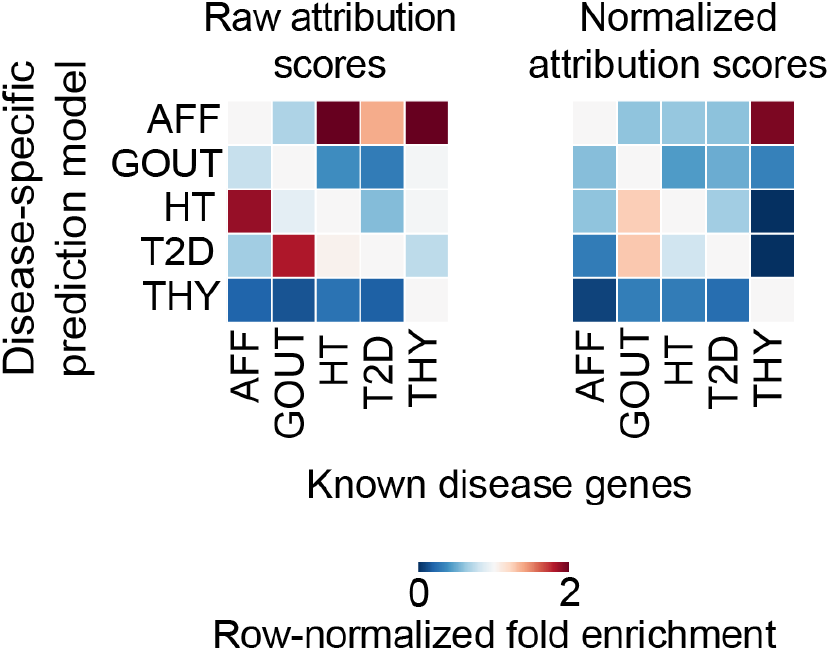
Normalized attribution scores lead to stronger disease-specific enrichment of disease associated genes. Heatmaps of row-normalized fold enrichment of known disease genes (columns) among the top attributed genes of each disease-specific FastBINN model (rows) for five traits. Each row is normalized across diseases so that values reflect the relative enrichment of each disease gene set within a given model. For a well specified model, enrichment is expected to concentrate on the diagonal, where a disease-specific model attributes importance to that disease’s known genes. Attribution score normalization sharpens this diagonal structure and reduces off-diagonal enrichment compared to raw scores, indicating that score normalization prioritizes disease-specific signal.

**Figure S4.**
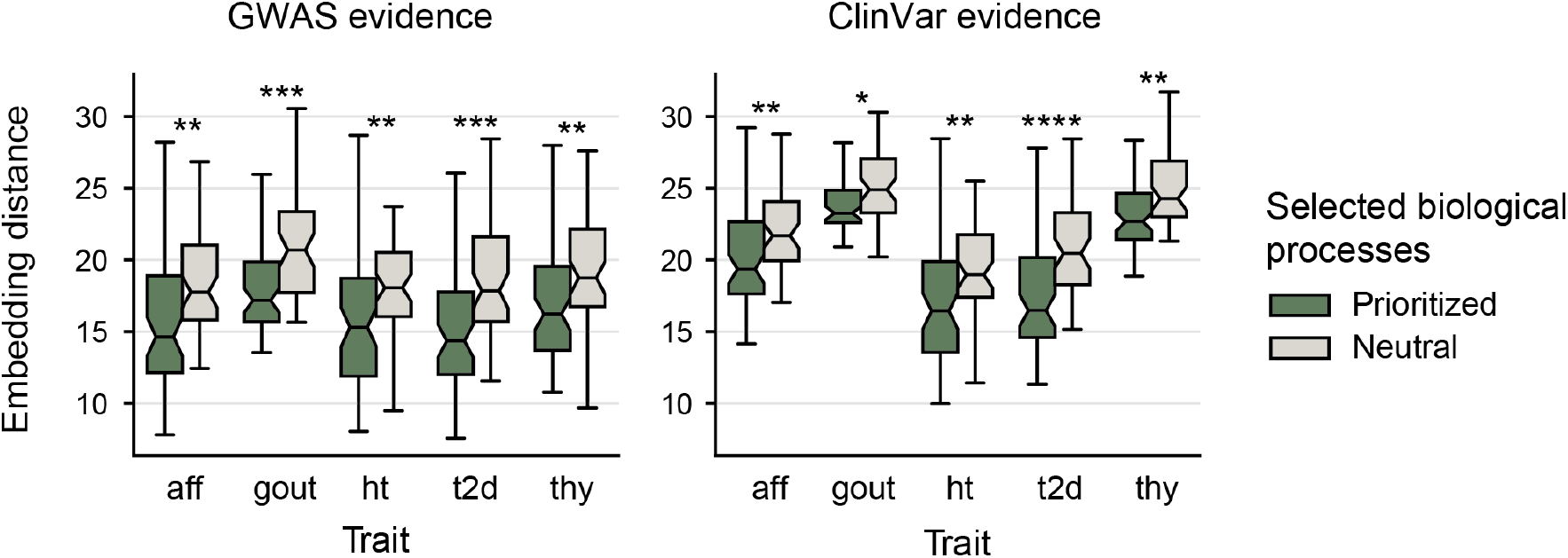
Independent validation of normalized FastBINN attributions by trait using a latent-neighborhood disease gene prioritization framework. Per-trait version of the collapsed analysis shown in Fig. 2e. For each trait, biological processes prioritized by FastBINN are closer in embedding space to disease representations from Open Targets (GWAS) and the European Variation Archive (ClinVar) than size-matched sets of mid-ranked biological processes sampled around the 50th percentile of the normalized attribution scores, supporting the biological relevance of normalized attributions beyond direct GWAS overlap.

**Figure S5.**
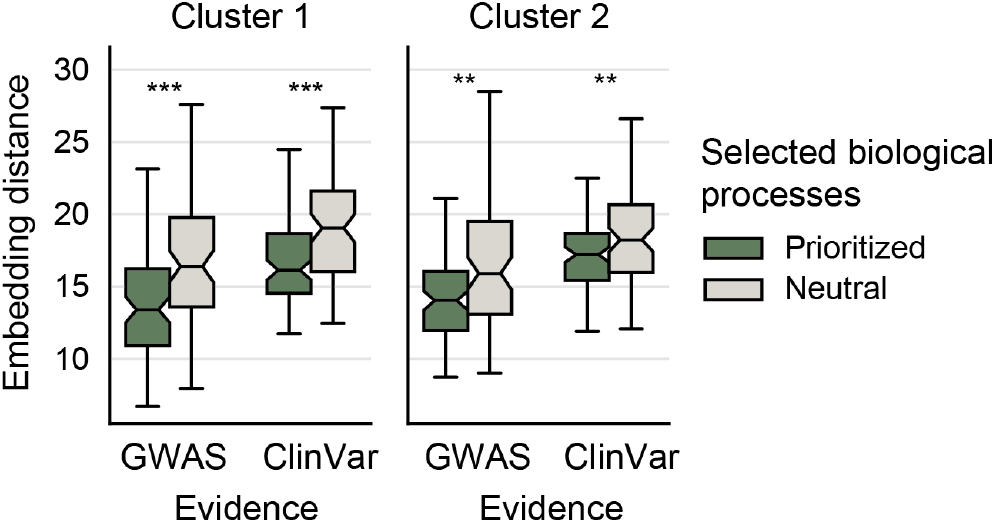
Independent validation of normalized FastBINN attributions by cluster using a latent-neighborhood disease gene prioritization framework. Per-cluster version of the collapsed analysis shown in Fig. 4a. For each cluster, biological processes prioritized by FastBINN are closer in embedding space to disease representations from Open Targets (GWAS) and the European Variation Archive (ClinVar) than size-matched sets of mid-ranked biological processes sampled around the 50th percentile of the normalized attribution scores. This supports the biological relevance of prioritized biological processes in both clusters.

**Figure S6.**
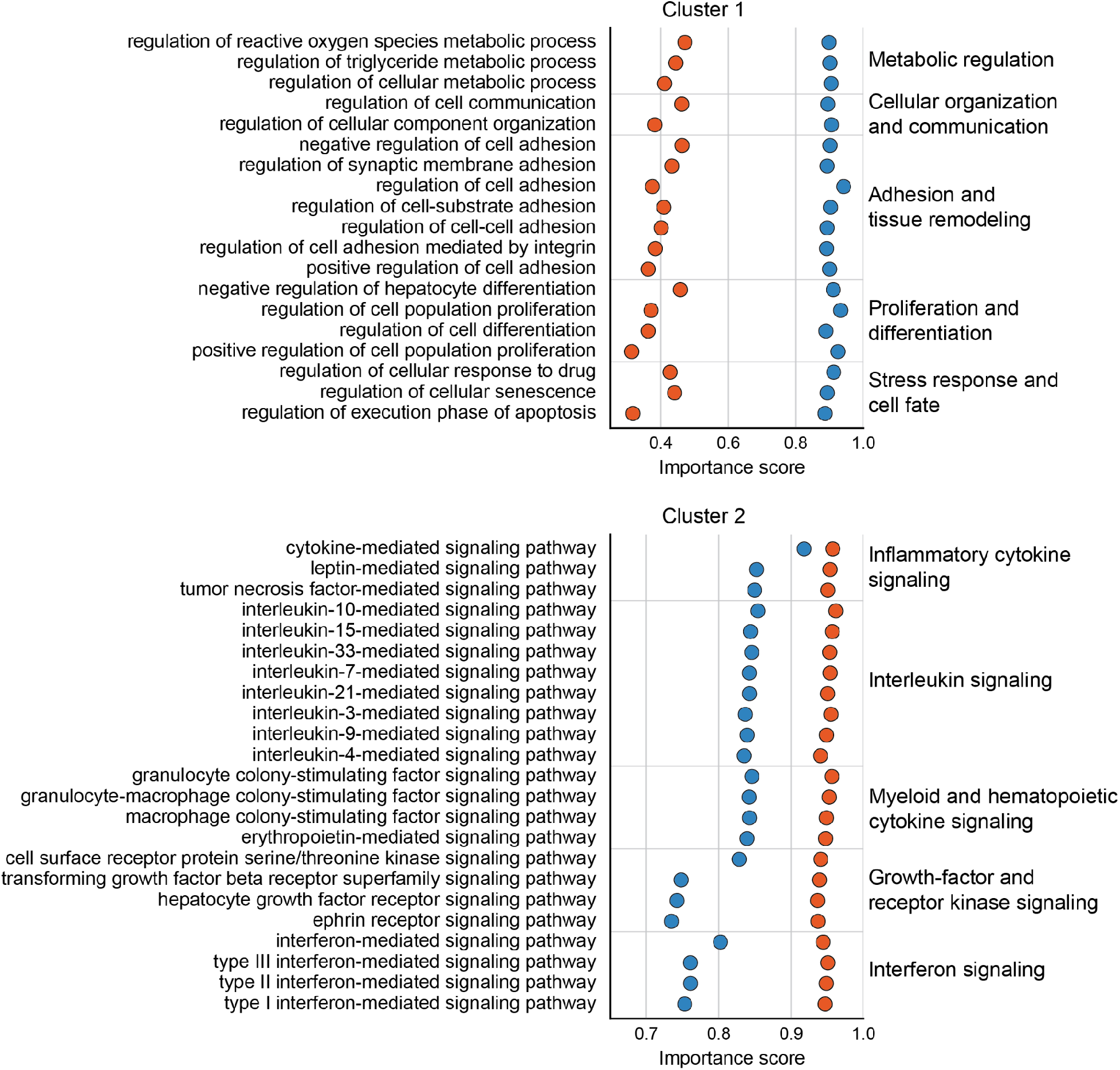
Full set of biological processes preferentially prioritized by the two main model clusters. Expanded version of Fig. 4c, showing all biological processes mapped to broader biological processes. The first cluster emphasizes metabolism and inflammation, the second one focuses on interleukin and cytokine signalling.

